# The change of lymphocyte subsets and inflammatory cytokine in BK viruria

**DOI:** 10.1101/2023.11.07.566078

**Authors:** Yu Huan Jiang, Lei Yuan, Qiang Chen, Hui Min He, Yang Liu, Lan Min Lai

## Abstract

**Background:** Polyomavirus BK (BKV) infection as a serious complication after kidney transplantation. The process of infections in kidney transplant recipients is viruria, viremia, and BKVAN. The difference between BK negative and BK viruria in kidney recipients has not been defined.

**Patients and Methods:** We compared post-transplant lymphocyte subsets 、blood cytokines、urine cytokines levels of 19 renal transplant outpatients with (BK-positive) or without BK viruria (BK-negative, n=20), and 20 healthy controls (HCs). Group of BK-positive divide into low-(n=4) and high-level (n=15) According to BK viral load (VL).Immune cells including T cells, B cells, and natural killer (NK) cells、interleukin-2(IL-2),IL-5,IL-6, IL-1β, IL-10, IL-8, IL-17A,IL-4,IL-12P70, interferon-α(IFN-α), IFN-γ,and tumor necrosis factor-α (TNF-α)were determined by flow cytometry.

**Results:** BK-positive patients showed higher urine IL-1β (*P*=0.040), IL-10 (*P=* 0.010), IFN-γ (*P* =0.002), and TNF-α (*P* =0.027) than BK-negative patients. Compared with HCs, BK-negative patients had lower urine IL-1β (*P*=0.04), IL-10 (*P=*0.01), TNF-α (*P* =0.027) and IFN-γ (*P* =0.004),suggesting that cytokine expression regulation BK-infection.

**Conclusion:** BK-positive renal transplant recipients, especially those with high VL, showed strong inflammatory cytokine responses with increases of urine IL-1β, IL-10, IFN-γ, and TNF-α. Our data suggest that monocyte- and Th-2-induced cytokines are involved in the pathogenesis of BKV-associated nephropathy.

## Introduction

BK virus was first discovered in a kidney transplant recipient who presented with a ureteral stricture in 1971[1].However, it was only in 1993 that the first definitive biopsy-proven case of BKVAN was described[2].BK virus infection is common in the general population, with seroprevalence rates of over 90% by 4 years of age[3, 4].The primary routes for transmission of the virus are from mucosal contact including the oral, gastrointestinal, and respiratory tract[5].Then, BK virus remains dormant in the kidney and uroepithelial cells resulting in lifelong latent/persistent infection. In immunosuppressed individuals,BK virus reactivation may cause kidney (BK-associated nephropathy) or bladder (hemorrhagic cystitis and ureteral stenosis) injury[6].The infection occurs in the following chronological stages—viruria, viremia, and allograft nephropathy[7, 8] .Viruria and viremia are detected in approximately 30% and 12% of kidney transplant recipients, respectively[9, 10] .Measuring BKV DNA in urine and serum is a useful and noninvasive tool for early detection and monitoring[11]. urine BK viral loads >8 log10 c/mL predict the onset of viremia, while plasma BK viral loads >4 log10 c/mL are associated with higher rates of biopsy-proven BKVAN[12-14] .Antiviral immunoregulatory markers like Gamma interferon (IFN-γ) might also affect the polyomavirus BK pathogenesis for its role in antiviral host defense, graft rejection, and regulative of the adaptive immune responses.Recent reports have shown that CD4+T cells likely have a direct role in controlling BKV infection, mediated through the expression of proinflammatory cytokines, including IFN-γ and tumor necrosis factor (TNF), and via the expression of the cytolytic molecule granzyme B[15].Analysis of the results showed that IFN-γ (rs12369470) CC genotype was significantly associated with susceptibility to BKV infection,it means Polymorphisms in the IFN-γ gene may confer certain protection or predisposition for BKV infection[16].Currently, the commonly used detection method includes detection of decoy cells in urine at cytological level, detection of DNA in urine and blood with polymerase chain reaction (PCR) and immunohistochemical examination[17] .Thus we want to find a new way to make an early diagnosis for BKVN with a rapid and effective detection method,as well, this method can intuitively reflect the severity of renal inflammation, highlighting the need for early intervention and therapy in clinical practice.

## PATIENTS AND METHODS

### Patients

We studied _BK viruria_、 cytokine responses in urine and blood、immune cells in blood of kidney transplant recipients between December 2021 and March 2023. Altogether, 171 patients were tested. Fifty-three patients (30.99%) were BK viruria positive (mean±SD 6.66±1.71 log_10_ copies/mL; range3.47-10.55 log_10_ copies/mL). For this study, we selected of the 19 BK viruria positive patients .The patients were treated with an immunosuppressive regimen consisting of triple drug therapy with calcineurin inhibitor (methylprednisolone, tacrolimus, enteric-coated mycophenolate sodium). They had no occurred acute rejection, anti-rejection therapy, or acute tubular necrosis during the last 3 months, and a lack of systemic or local signs of another acute viral or bacterial infection. The 19 BK-positive patients were compared with 20 of the 118 BK-negative patients who were similar in age, gender, immunosuppressive drugs, serum creatinine(Table 1). Four BK-positive patients with low VL (4.24 –5.43log_10_) and fifty with high VL (6.12–10.12 log_10_). BK-positive patients had consistent or periodic viruria, whereas BK-negative patients had BKV-negative urine results in all post transplant investigations. Kidney function was stable in all patients. Twenty healthy blood donors were tested as controls. Demographic data of BK-positive patients, BK-negative patients,and healthy controls (HCs) are shown in Table 1.

**TABLE 1.** Demographic data of healthy controls, BK-positive, and BK-negative patients.All data are given as mean±SD. P values were calculated using χ2, Fisher’s exact, and Mann-Whitney U test. MP, methylprednisolone; Tac, tacrolimus; EC-MPS, enteric-coated mycophenolate sodium.

| Parameter | BK-positive(n=19) | BK-negative (=20) | P | Healthy controls(n=20) |
| --- | --- | --- | --- | --- |
| Age (yr;(x±SD)) | 41.26±9 | 38.65±10 | 0.673 | 35.6±12.58 |
| Female (n) | 7 | 6 | 1.000 | 8 |
| Urine BK viral load (log10) | 7.29±1.64 | - | <0.001 | - |
| Serum Cr (mg/dL) | 147.4±88.1 | 164.7±163.3 | 0.950 | 74.32±10.46 |
| Immunosuppressive drugs |  |  |  |  |
| EC-MPS(n) | 9 | 9 | 1.000 | - |
| MP dose(n) | 6 | 13 | 0.227 | - |
| Tac (n) | 17 | 19 | 1.000 | - |
| PrednisoloneAcetate(n) | 13 | 17 | 0.952 | - |
| EC-MPS(mg/mL) | 340±101.98 | 320±74.83 | 1.000 | - |
| MP dose(mg/mL) | 306.67±203.52 | 220±221.46 | 0.868 | - |
| Tac (ng/mL) | 7.05±1.98 | 6.81±1.44 | 0.393 | - |
| PrednisoloneAcetate(mg/mL) | 13.85±11.46 | 13.18±5.055 | 0.953 | - |
| Post-Tx days of investigation(mean±SD) | 384.53±290.81 | 401.5±247.73 | 0.802 | - |

### Healthy Individuals

In samples from 20 healthy lab staff members, cytokine levels were assessed. None of the individuals had any acute or chronic illnesses, a urinary tract infection, or were taking any medications. In controls, urine real-time PCR was used to determine BKV and urine cultures were used to detect bacteria. Eight women were menstrually free at the time of the inquiry. A questionnaire was used to collect anamnestic data.

## Methods

### Isolation of BKV nucleic acid and amplification by real-time PCR

First, 100μl blood was poured into 1.5 ml centrifuge tube and then added with 50 μl concentrated solution. After being shaken up for 15 s, the tube was centrifuged in a table centrifuge at 13,000 r/min for 10 minutes. 150μl supernate was absorbed from the upper layer and abandoned. Afterwards, 25 μl lysis solutions were added into centrifuge tube. Sediment left in the centrifuge tube was removed out with sucker and then was blew and washed for five times. After the sediment was thoroughly scattered through 15 shaking, it was incubated at 100 °C. Ten minutes later, it was centrifuged at 13,000 r/minutes for 10 minutes. The supernate obtained, i.e., purified DNA solution, was transferred to new centrifuge tube. Urine loaded in tube was first shaken for 15 s. Then 1 to 1.5 ml urine was put into centrifuge tube and centrifuged at 13, 000 r/min for 10 min. After the supernate was removed, 50 ul lysis solution was added. Then sediment left was removed out from the centrifuge tube with sucker and blew and washed for 5 times as well. The following steps were the same as blood specimen.5μl BKV negative quality control products, specimen and quantitative BKV standard substance I ∼ IV were taken and added into PCR tubes. Then they were centrifuged at 2, 000 r/min for 15 s to throwing the liquid on the wall of tube to the bottom of tube. If bubbles flicked the wall of tube, the liquid was centrifuged once more. Then PCR amplification was performed on a ABI7500 instrument (ThermoFisher Scientific, Waltham, MA))with a thermocycling profile at 37°C for 2 min,95°C for 3 min followed by 40 cycles at 94°C for 15 sec, and at 60°C for 35 sec. The detection limit was determined at 2000 copies/mL[17].

### Lymphocyte Subset Detection

Peripheral venous blood samples (EDTA anticoagulated) were collected from all participants. The absolute numbers and percentages of CD3+ T cells, CD4+ T cells, CD8+ T cells, B cells, and NK cells were determined using a 6-color TBNK Reagent Kit (antiCD45-PerCP-Cy5.5 (2D1), anti-CD3-FITC (UCHT1), anti-CD4-PC7 (RPA-T4), anti-CD8-APC-Cy7 (H1T8a), anti-CD19-APC (H1B19), anti-CD16-PE (CB16), and anti-CD56-PE (MEM-188)) (QuantoBio Technology, Beijing, China) with QB cell-count tubes, according to the manufacturer’s instructions. The kit uses a lyse-no-wash staining procedure and provides absolute cell numbers. At 37 °C, 50 μL of whole blood was stained with 20 μL of a 6-color TBNK antibody cocktail for 15 min. After adding 450 μL of RBC lysis solution (QuantoBio Technology, Beijing, China) and after 15 min of incubation, the samples were analysed using a DxFLEX flow cytometer (Beckman Coulter, Fullerton, CA, USA). All data were analysed using FlowJo software (version 10.5.3, FlowJo LLC, Ashland, OR, USA).

### Lymphocyte phenotype analysis

Peripheral venous blood samples (EDTA anticoagulated) was collected from study participants. The following monoclonal antibodies were added to 100 μl of whole blood: anti-CD45, anti-CD3, anti-CD4, anti-CD8, anti-CD28, anti-HLA-DR(Beckman Coulter, Villepinte, France). Isotype controls with irrelevant specificities were included as negative controls. All of these cell suspensions were incubated for 20 min at room temperature. After lysing red blood cells, the cells were washed and resuspended in 200 μl of PBS. The cells were then analyzed with DxFLEX flow cytometer (Beckman Coulter, Fullerton, CA, USA).

### Cytokine Profile Analysis

Serum and urine samples were collected from all participants. The blood was centrifuged at 1000g at 20 °Cfor 20 min. The serum was carefully harvested and determined for the Th1/Th2 cytokines by flow cytometry immediately or the serum was temporarily stored at 2–8 C until analysis if the situation was not so urgent (usually within 12 h). Concentrations of IL-1-β,IL-2, IL-4, IL-5,IL-6, IL-8,IL-10,IL-12p70,IL-17A, TNF-α and interferon (IFN)-α were quantitatively determined by BD™ CBA Human Th1/Th2 Cytokine Kit II (BD Biosciences, San Jose, CA) as described previously[18]. The minimum and maximum limits of detection for all 12 cytokines were 1.0 pg/mL and 5000 pg/mL, respectively.

### Statistical Analysis

A statistical analysis was performed using SPSS version 23.0 and GraphPad Software 9.0. Continuous variables are expressed as mean ± SEM. The comparisons were performed using the χ2or Fisher’s exact test for categorical variables and using the Mann–Whitney U test(for two groups) or Kruskal–Wallis H test (for three groups) for continuous variables. Prognostic factor analysis was performed by univariate analysis using log-rank test. Multivariate analyses were performed in logistic regressionand odds ratio(OR) was calculated, respectively. Receiver operating characteristics (ROC) curves were performed to evaluate the prediction power of IFN-γ.All p-values were two-tailed, and differences with p < 0.05 were considered statistically significant.

## RESULTS

### Development of BK viruria and BK infection

thirty-nine of nineteen patients (49%) developed BK viruria ranging from 1.7×10^4^– 1.3×10^10^ copies/mL of urine (median 3.3×10^7^ copies). The cumulative incidence of BK viruria peaked between two months and a year, when more than 52.4% of individuals were confirmed to have the condition.The incidence of BK viruria at various time intervals during follow-up is shown in Table 2.

**Table 2.** Incidence of BK viremia at different time-points.

| Time point | Tested(n) | Positive(n,%) |
| --- | --- | --- |
| 8 weeks | 4 | 4(100%) |
| 6 months | 8 | 3(37.5%) |
| 1 year | 6 | 4(66.7%) |
| 2 years | 11 | 5(45.5%) |
| Uper 2 year | 8 | 5(62.5%) |

Urine levels of cytokines were assessed in 39 transplant recipients. The results are shown in Table 3.

**Table 3.**
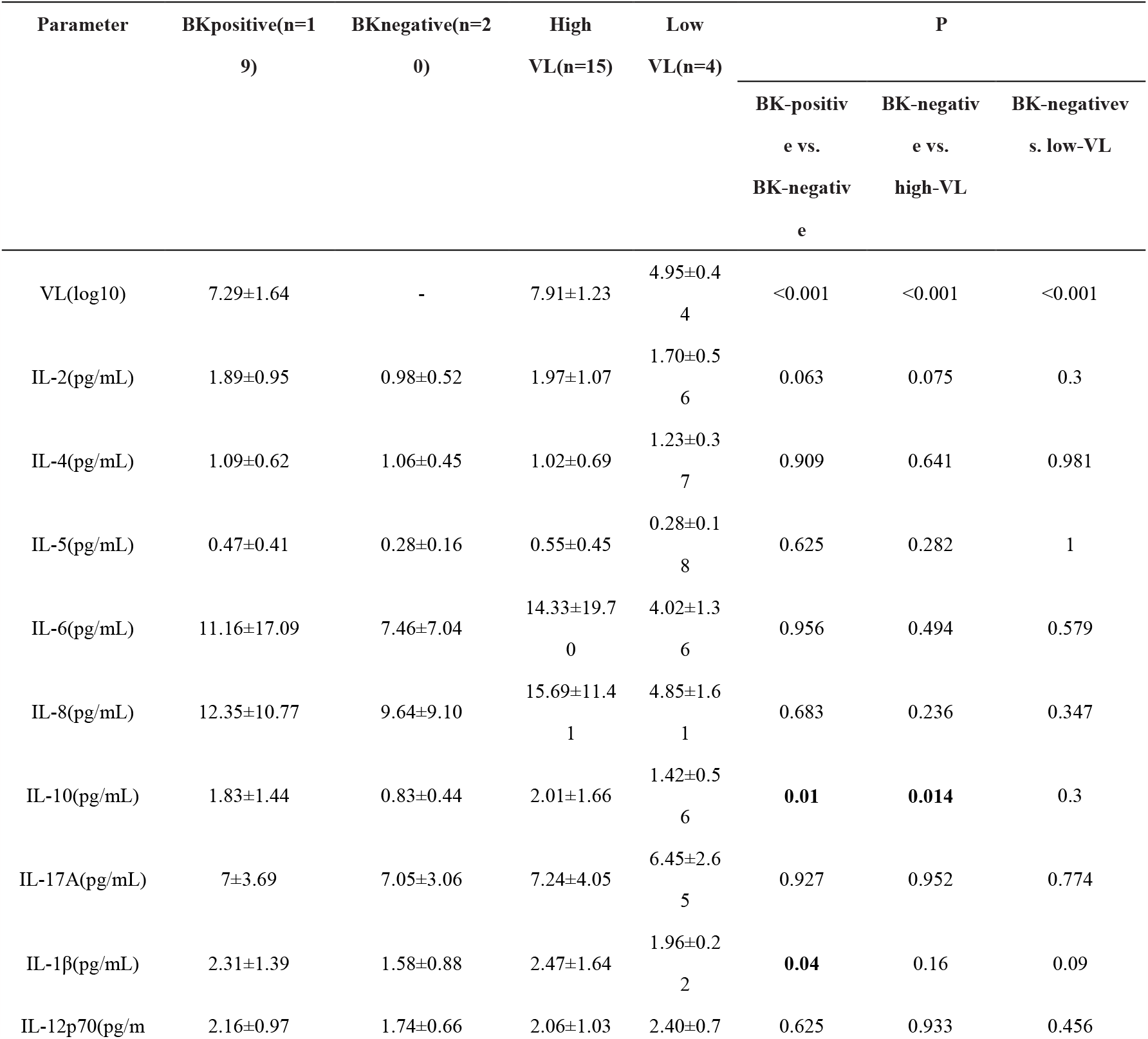
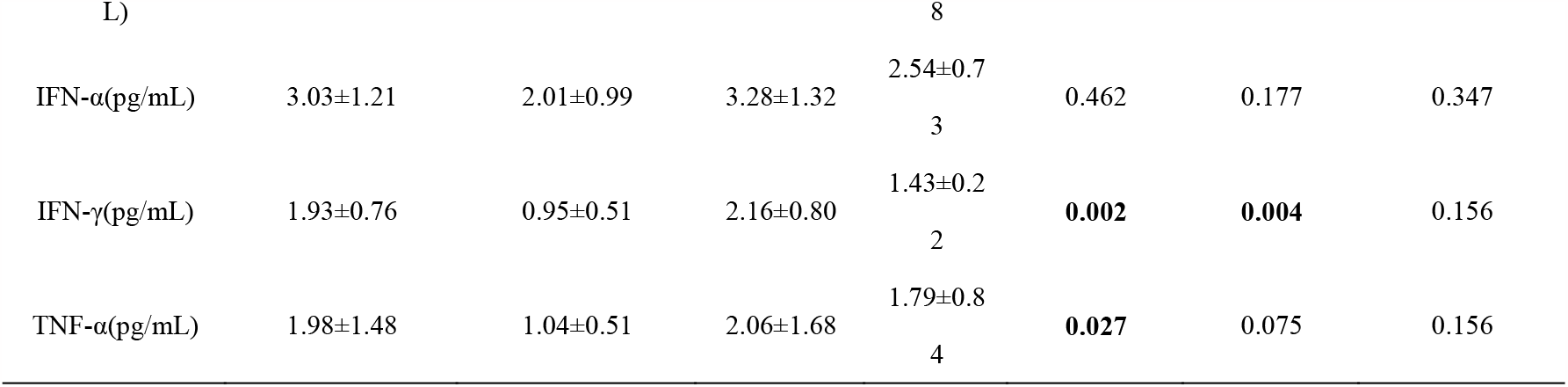
Urine levels of cytokines were assessed in 39 transplant recipient*s*.*P* values were calculated using the Wilcoxon test. Adjustment for multiple testing was performed according to the method of Bonferroni. Only *P* values of 0.05 after adjustment were considered to be significant and printed bold in the tables. *a* Values are represented as mean±SD. VL, viral load; HCs, healthy controls;IL, interleukin;TNF, tumor necrosis factor; IFN, interferon.

### IL-10

Urine IL-10 was high in BK-positive transplant recipients than in BK negative patients (1.83±1.44 vs. 0.83±0.44 pg/mL, P=0.01). However,patients with high VL had significantly higher IL-10 than BK-negative patients, suggesting upregulation of urinary IL-10 in the subset of recipients with BK positive,especially high VL .(Table 3, Fig. 1A).

**Fig 1.**
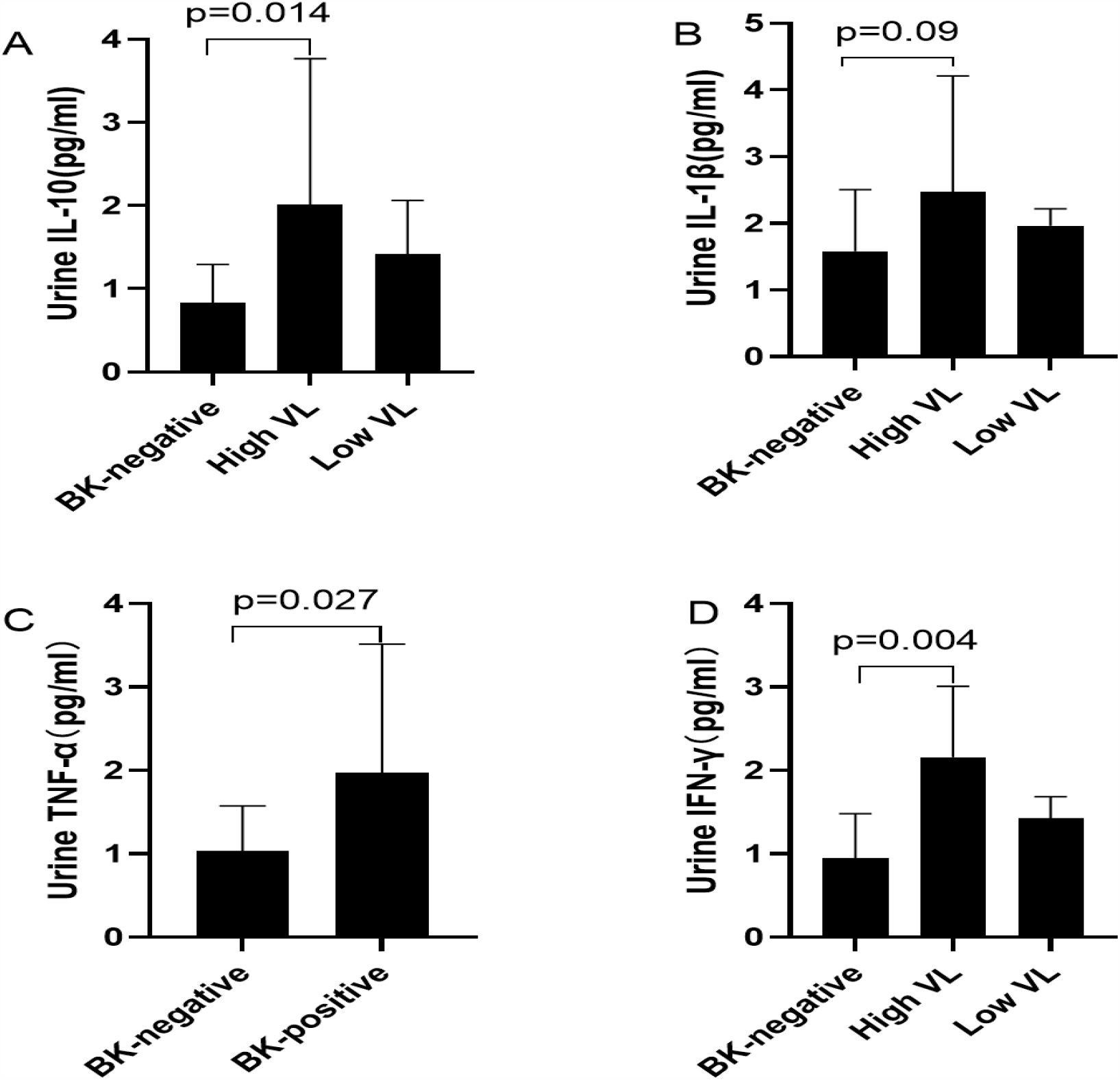
Urine IL-10、IL-1β、TNF-α、IFN-γ levels in renal transplant recipients with high BK viral load (n=15), low BK viral load (n=4), without viruria(n=20).

### IL-1β

Urine IL-1β level was higher in BK-positive transplant recipients than in patients without viruria(2.31±1.39 vs. 1.58±0.88 pg/mL, P= 0.04). Patients with low VL had higher urine IL-1β levels than patients without viruria too. suggesting an increase of urine IL-1β with viral infection (Table 3, Fig. 1B).

### TNF-α

Urine TNF-α was significantly higher in BK-positive than in BK negative patients (P=0.027; Table 3, Fig. 1C).

### IFN-γ

Urine IFN-γ was higher in BK-positive patients (1.93±0.76 vs. 0.95±0.51 pg/mL, P 0.002), high VL (2.16±0.8 vs 0.95±0.51pg/mL, P 0.004) than BK-negative recipients, suggesting increased urinary IFN-γ in BK-positive transplant recipients (Table 3, Fig. 1D).

Blood levels of cytokines and immunocytes were assessed in 39 transplant recipients. The results are shown in Table 4.

Blood IL-2 level was higher in BK-positive(1.93±3.1vs.0.15±0.37,P=0.001) and BK-negative (0.73±0.41vs.0.15±0.37,P<0.001)than in HCs;IL-17was significantly higher in BK-positive(7.29±3.55vs.4.3±0.51,P=0.001) with BK-negative (5.91±3.83vs.4.3±0.51,P=0.012) patients than in HCs . According to the former study, it showed that blood IL-4 was higher in HCs than in BK-positive(0.67± 0.82vs.1.8±0.37,P<0.001) and BK-negative(0.3±0.22vs.1.8±0.37,P<0.001) .In comparison to BK-positive (2.66±2.82vs.4.39±0.63,P<0.001) and BK-negative (2.12±1.07vs.4.39±0.63,P<0.001) patients, HCs had greater IL-10 levels.IFN-γ levels were higher in HCs than they were in BK-positive (1.58±1.11vs.3.65±0.34,P <0.001) or BK-negative (1.36±0.53 vs.3.65±0.34,P<0.001).Blood CD3% numbers were more substantial in BK-positive (77.29±12.59 vs. 68.27±8.33, P=0.001) and BK-negative (79.6±9.63vs.68.27±8.33,P<0.001) subjects than in HCs.Blood CD8% counts in BK-positive (33.0611.08vs.26.056.21, P=0.025) and BK-negative (32.588.36vs.26.056.21, P=0.043) individuals were higher compared to in healthy controls.HCs had deeper B cell counts than BK-positive (92.53±87.19vs.277.55±113.17,P<0.001) or BK-negative (151.07±124.57 vs.277.55±113.17, P=0.003) individuals.B% levels were heavier in HCs than they were in BK-positive (7.55± 4.85 vs. 14.92 ±6.92, P=0.001) or BK-negative (6.49±2.87 vs.14.92±6.92,P<0.001).

### ROC-Curve Analysis

ROC-curve analysis was used to compare urine cytokines of a certain patient group with those of other patient groups. ROC-curve analysis showed high sensitivity and specificity of urine IL-10 and IFN-γ for the identification of patients with BK-positive patients. Sensitivity and specificity for urine IL-10, calculated with a cutoff level of 1.11 pg/mL, were 83.3% and 81.8%, respectively (area under the curve 0.832, 95% confidence interval 0.657–1.000, P=0.006). The corresponding values for urine IFN-γ, cutoff level of 1.380 pg/mL, were 84.6% and 90.9%, respectively (area under the curve 0.902, 95% confidence interval 0.641 –0.989, P 0.009; Fig. 2).

**Fig2:**
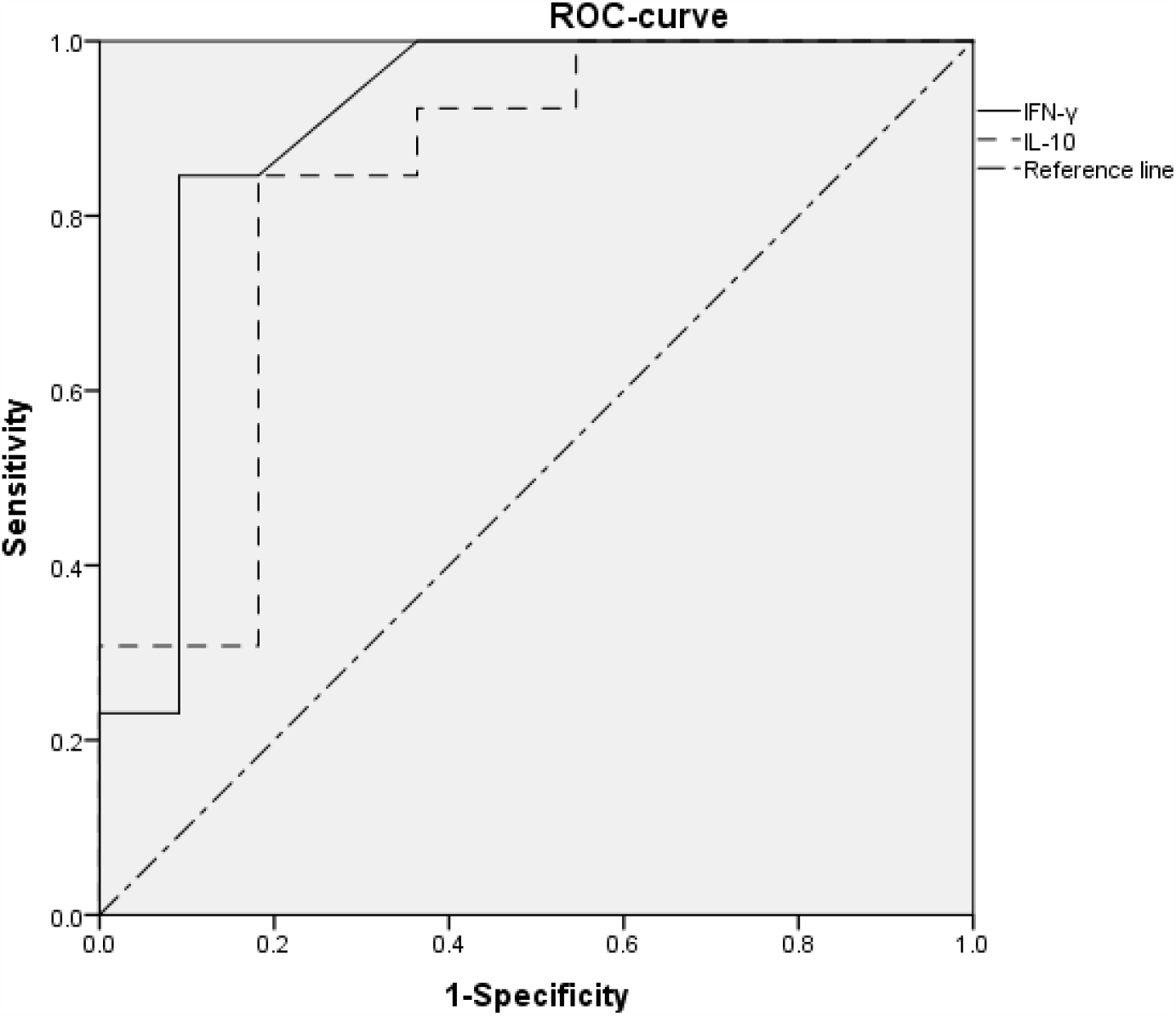
ROC analysis demonstrated a significant relationship of urine IL-10 and IFN-γ with BKV level in urine.

Area under the curve 0.832and 0.902, respectively, suggesting that high urine IL-10 (cutoff value >1.11 pg/mL) and urines IFN-γ (cutoff value >1.38 pg/mL)are related to urine BKV level.Sensitivity was 83.3% and 84.6%, respectively, and specificity was81.8% and 90.9%, respectively.

### Risk factors analysis for BK-positive patients

Multivariate analysis revealed that IFN-γ(OR 0.022,95%Cl 0.001-0.499 P=0.016) is the only one protective factor for BK virus infection.

### CD4^+^ T cell count negative Correlated with BK VL in BK-positive patients

There were significant negative correlations of urine BK VL with Blood CD4^+^ T cell count in BK-positive patients(r=-0.529,P=0.02; Fig 3).

**Fig 3:**
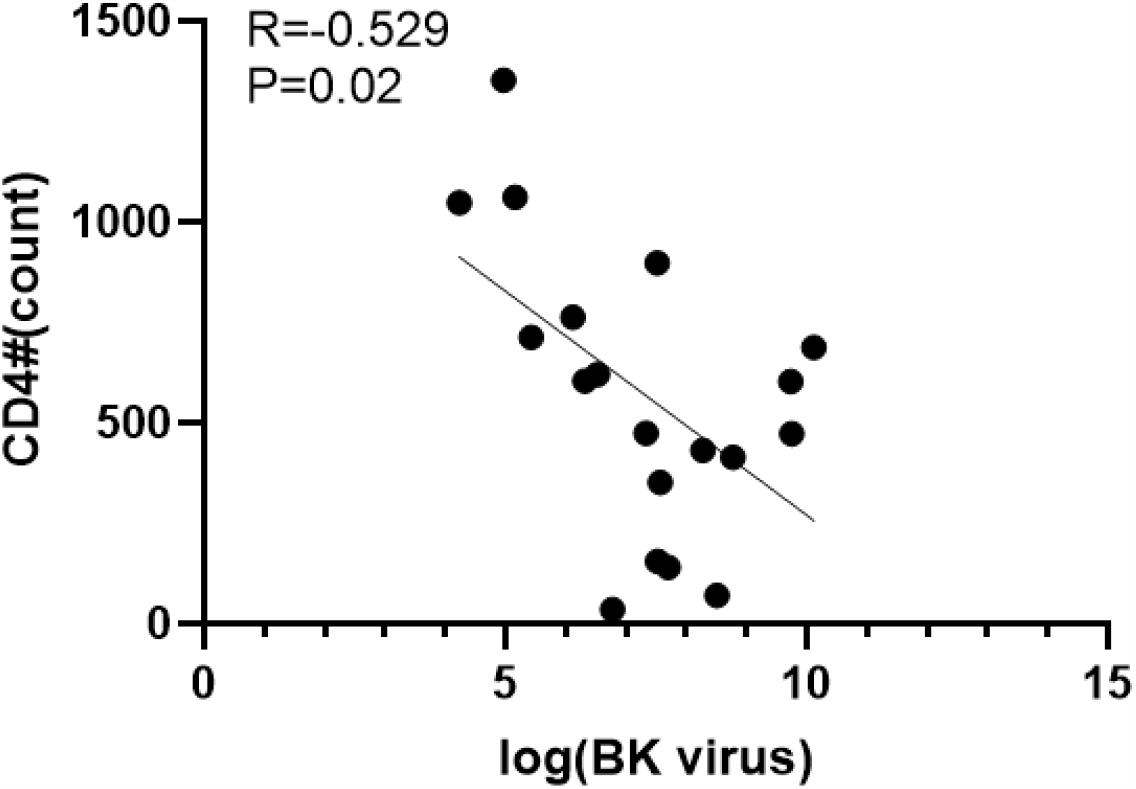
Significant positive correlations were found between levels of CD4 T cell count and the load of BK virus in BK-positive patients.

### TNF-αlevel is negative Correlated with BK VL in patients with low VL and high VL

In patients with low VL, there were significant negative associations between urine BK VL and TNF levels (r=-0.941, P=0.002;Fig4A).However, there is a lack of correlation between BK viral load and TNF-in BK high VL patients(r=0.229, P=0.523;Fig4 B).

**Fig 4:**
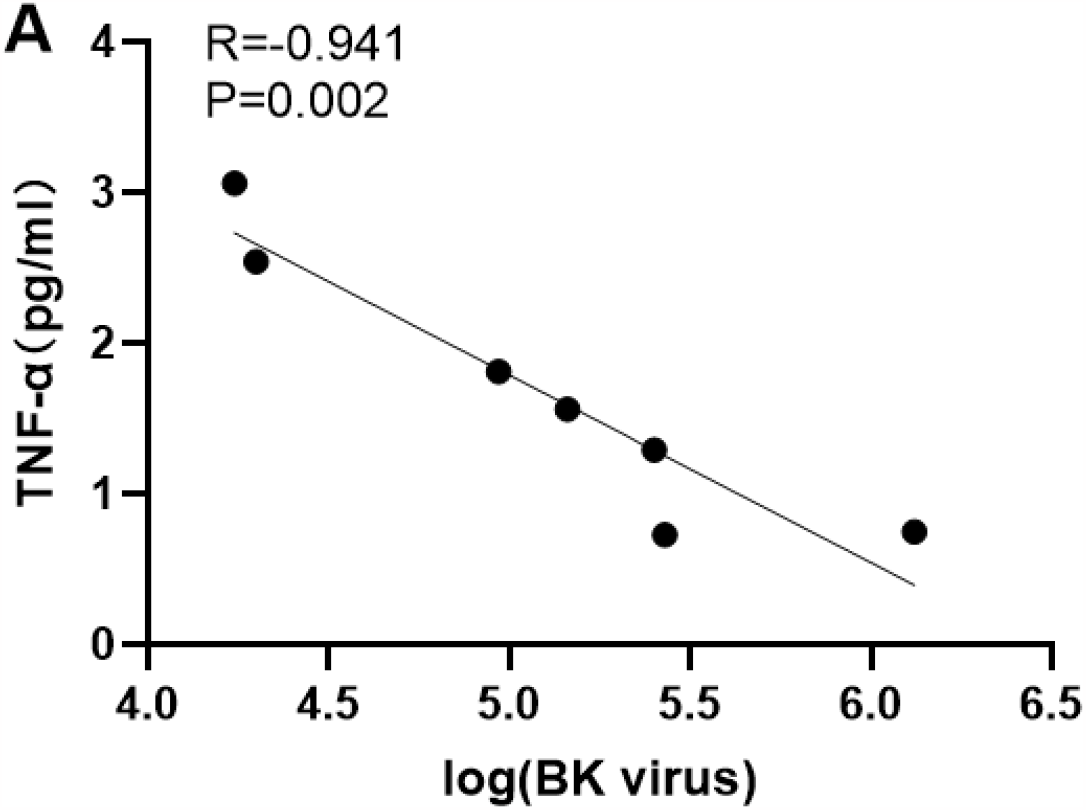
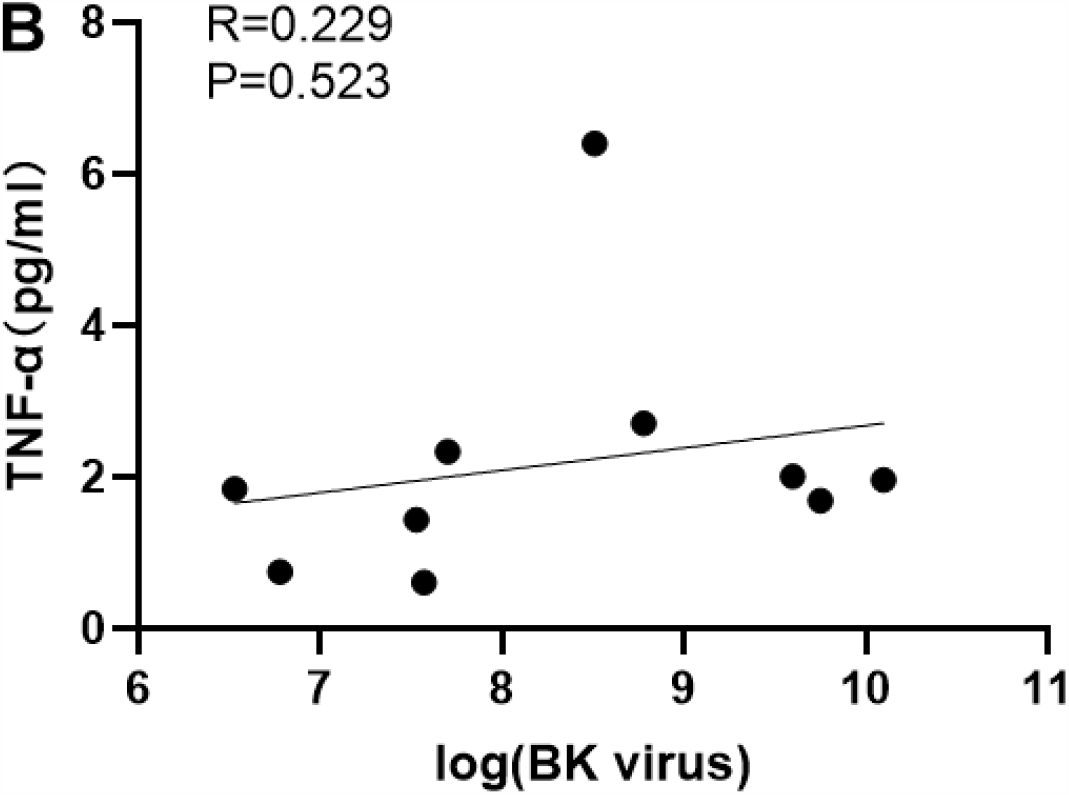
Significant positive correlations were found between levels of urine TNF-α and the load of BK virus in low VL patients (Fig 4A),the connections between urine TNF-α levels and BK virus load in patients with high VL(Fig 4B).

## DISCUSSION

A symptom of active replication of the virus in transplant recipients is excretion of the virus in the urine, i.e., viruria (80% of patients), while viraemia, where virions are detected in the blood, appears in 10–15%, and severe tubule interstitial nephritis (BKPyVAN) about 8%[19].In some patients with BKPyVAN irreversible kidney damage takes place, resulting in loss of the transplant[20, 21].The site of BK virus persistence is the kidneys,so testing lymphocytes and cytokines in blood or urine can provide a more accurate reflection of the state of local and systemic inflammation.Currently, real-time fluorescence quantitative PCR is advised for detecting the DNA load of BK virus. By comparing the laboratory results of BK positive and BK negative patients, we expect to discovry a novel indicator to detect early BK virus infection.We discovered that IL-10、IL-1β、IFN-γ and TNF-α were greatly elevated in BK positive patients when comparing the urine cytokines of BK positive and BK negative patients.

Our study demonstrates a substantial inverse relationship between the BK viral load and the absolute number of CD4+T cells in patients who test positive for the BK virus. In patients with BKVN nephropathy, we discovered through a study of the literature that there is a negative association between the amount of BKV-DNA and the BKV-specific CD4+T cell response[22]. Patients with bone marrow transplants exhibit a negative connection between BK virus reactivation and CD4+T cell count, which is very congruent with the findings of our study[23].

Our results show TNF-α level was significantly negatively correlated with BK Viral load, furthermore TNF-α was found through literature review it can inhibit the replication of BK virus[24], which is consistent with our research results. In our study, The level of TNF -α was negatively correlated with the concentration of BK virus during low viral load, however at high BK virus load the concentration of TNF-αis already very low, so it cannot inhibit the replication of BK virus, therefore TNF-α concentration has no obvious correlation in high BK Viral load.

IL-10 targets both innate and adaptive immune responses .In particular during the resolution phase of infection and inflammation, IL-10 exerts immunosuppressive activities to prevent tissue damage produced by excessive and uncontrolled inflammatory effector responses and to maintain equilibrium to gut microorganisms.In certain situations, IL-10 expression is helpful in the eradication of viruses .Coproduction of IL-10 and IFN-γ during influenza infections promotes the buildup of anti-influenza antibodies in the lung mucosa[25]. The fact that persistent infections with HCV, HBV, HIV, and LCMV have been linked to this loss of T cell function raises the possibility that a common immunosuppressive mechanism may inhibit T cell activity. These methods result in the induction of inhibitory receptors [26],or increased levels of IL-10 in the body[27, 28].

A powerful inflammatory cytokine called IL-1β is responsible for recruiting immunological and inflammatory cells to the site of an infection or injury. The development of adaptive immune responses is influenced by IL-1β[29].Both acute and chronic inflammation caused by viral infections respond to IL-1β[30, 31]. It is created in response to motifs inside macromolecules that accumulate during microbial infection known as pathogen-associated molecular patterns (PAMPs)[32, 33].According to preliminary experimental results, the level of IL-1β is higher in BK positive patients than in BK negative individuals.Therefore, we speculate whether the BK virus promotes IL-1β secretion through the production of inflammasomes.

Interferons (IFNs) are important antiviral cytokines which act as the body’s initial line of defense against infection[34]. IFN-gamma, a cytokine that promotes inflammation, is the sole member of the type II interferon family. During the adaptive immune response, it is made by T-cells and innate immune cells, including natural killer (NK) and antigen-presenting cells.According to Tony Fiore’s research, IFN-inhibits the replication of the BK virus by activating the Jak Stat signaling system[35].Using real-time fluorescence PCR technique and the MRNA levels of kidney transplant recipients, Neda Zarei compared the detection of IFN in BK-infected and uninfected patients with healthy controls. IFN-γ Patients with BK infection had significantly higher levels of mRNA expression than uninfected patients and healthy comparison groups[36].IFN- (rs12369470) CC genotype is strongly related with BKV infection susceptibility, while IFN- +874 (rs2435061) TT and (rs2406918) CC genotypes appear to be markers for protection against BKV infection, according to research done by Vu and Don on 251 kidney transplant patients. Therefore, IFN-γ gene polymorphisms may confer a degree of protection or propensity to BKV infection[37].Our findings demonstrate that BK-positive patients’ urine IFN-γ levels were obviously more than patients with BK negative . IFN-γ has great sensitivity and specificity at 1.38 pg/ml, according to the examination of receiver operator characteristics.Multiple factor analysis shows IFN-γ is a protective factor against BK virus infection, which is consistent with the findings of other studies.

The fact that there was no discernible difference in the number and percentage of blood cells between BK positive and negative patients. Until we analyzed the blood cells of both groups indicate the infection was mostly located in the urinary system and did not spread into the blood.Fascinatingly, patients with low-level VL exhibited similar reactions to those of individuals with BK negativity.According to our data, BK viruria of less than 7 log10 copies/mL is not important for therapy consideration, as suggested by the literature, and the inflammatory response in kidney transplant recipients during BKV replication is closely correlated with VL levels. It may be useful to monitor urine cytokines, particularly IFN-γ, to aid in the diagnosis of BKV infection and the ongoing care of patients who have BK viruria.Fewer patients have received kidney transplantation in our study due to a lack of kidney supplies, and our sample size is relatively modest due to clinicians’ poor understanding of the BK virus and kidney impairment. Furthermore,the BK virus Nucleic acid test at the hospital is relatively short, our observation period for patients who test positive for the virus is also shortly. Nevertheless, we will continue to monitor the follow-up results of these positive patients.

## Acknowledgments

The authors wish to thank the patient for joining in this study and all the staff members at our institution.

---

**Table 4.Blood levels of cytokines and immunocytes were assessed in 39 transplant recipients and 20 HCs.**

